# CD5L is a canonical component of circulatory IgM

**DOI:** 10.1101/2023.05.27.542462

**Authors:** Nienke Oskam, Maurits A. den Boer, Marie V. Lukassen, Pleuni Ooijevaar-de Heer, Tim S. Veth, Gerard van Mierlo, Szu-Hsueh Lai, Ninotska I.L. Derksen, Victor C. Yin, Marij Streutker, Vojtech Franc, Marta Siborova, Mirjam Damen, Dorien Kos, Arjan Barendregt, Albert Bondt, Johannes B. van Goudoever, Carla J.C. Haas, Piet C. Aerts, Remy M. Muts, Suzan H.M. Rooijakkers, Gestur Vidarsson, Theo Rispens, Albert J.R. Heck

**Affiliations:** Sanquin Research and Landsteiner Laboratory, Department of Immunopathology, Amsterdam UMC, Amsterdam, Netherlands; Biomolecular Mass Spectrometry and Proteomics, Bijvoet Center for Biomolecular Research and Utrecht Institute for Pharmaceutical Sciences, University of Utrecht, Padualaan 8, Utrecht 3584 CH, the Netherlands; Netherlands Proteomic Center, Padualaan 8, Utrecht 3584 CH, the Netherlands; Amsterdam UMC, Vrije Universiteit, University of Amsterdam, Emma Children’s Hospital, 1105 AZ Amsterdam, The Netherlands; Department of Medical Microbiology, University Medical Center Utrecht, Utrecht University, Utrecht, the Netherlands; Sanquin Research and Landsteiner Laboratory, Department of Experimental Immunohematology, Amsterdam UMC, Amsterdam, Netherlands

## Abstract

Immunoglobulin M (IgM) is an evolutionary conserved key component of humoral immunity, and the first antibody isotype to emerge during an immune response. IgM is a large (1 MDa), multimeric protein, for which both hexameric and pentameric structures have been described, the latter additionally containing a joining (J) chain. Using a combination of single-particle mass spectrometry and mass photometry, proteomics and immunochemical assays, we here demonstrate that circulatory (serum) IgM exclusively exists as a complex of J-chain-containing pentamers covalently bound to the small CD5 antigen-like (CD5L, 36 kDa) protein. In sharp contrast, secretory IgM in saliva and milk is principally devoid of CD5L. Unlike IgM itself, CD5L is not produced by B cells, implying that it associates with IgM in the extracellular space. We demonstrate that CD5L integration has functional implications, i.e., it diminishes IgM binding to two of its receptors, the FcαµR and the polymeric Immunoglobulin receptor (pIgR). On the other hand, binding to FcµR as well as complement activation via C1q seem unaffected by CD5L integration. Taken together, we redefine the composition of circulatory IgM as a J-chain containing pentamer, always in complex with CD5L.

## Introduction

Immunoglobulin M (IgM) is the first antibody isotype to emerge in ontology and during an immune response. IgM is integral to the initiation of the humoral immune response, but also for maintaining immune homeostasis through the induction of tolerance and for instance the clearance of apoptotic cells ^1, 2^. It is furthermore a potent activator of the classical pathway of the complement system and regulates antigen presentation and B-cell maturation through interactions with the IgM-specific receptors FcµR ^3^ and FcαµR ^4^. Additionally, secretory IgM plays an important role in mucosal immunity, as the integration of a small joining (J) chain allows it to be transported to mucosal surfaces via the polymeric immunoglobulin receptor (pIgR) ^5^. IgM is present throughout our body, with concentrations in serum, human milk and saliva of ca. 1500 mg/L ^6^, 2.8 mg/L ^7^ and 1.2 mg/L ^8^, respectively.

IgM is expressed by B cells as a precursory monomeric (H2L2), membrane-bound B cell receptor, which recognizes antigen and relays survival and proliferation signals for maturing B cells. When a B cell switches to IgM secretion through alternative splicing of the µ heavy chains (HC), it can concomitantly co-express the J-chain. Five IgM protomers, each consisting of two heavy chains coupled to two light chains, can combine with a J-chain to form one pentameric molecule, which has for decades been assumed to be the principal arrangement of IgM in circulation ^1, 9^. Furthermore, it has been shown that if IgM is expressed without J-chain it can assemble into hexamers ^10–12^. However, (monoclonal) hexameric IgM observed in circulation seems invariably linked to pathologies such as Waldenström macroglobulinemia or cold agglutinin disease ^10, 11^. Despite persistent speculation in literature, it is currently unclear what fraction, if any, of normal human circulatory IgM is hexameric ^12–14^.

Recently, several cryo-EM studies have shed more light on the detailed molecular structure of pentameric IgM. Rather than the previously assumed symmetrical pentameric arrangement, recombinant IgM with J-chain forms an asymmetrical pentamer resembling a hexameric structure, wherein the J-chain bridges the gap in place of a sixth IgM subunit ^15–19^. The core of these structures is comprised of an amyloid-like assembly of the J-chain with the C-terminal IgM tailpieces, which are responsible for this isotype’s propensity to oligomerize. On top of this assembly, the secretory component (SC) of pIgR can bind IgM through interaction with the J-chain and IgM heavy chains ^16, 19^.

Intriguingly, by using *in vitro* reconstitution, using negative stain EM, it was reported that a protein called CD5-Like molecule (CD5L) can also associate into the gap of murine J-chain-linked pentameric IgM and form a covalent attachment through a disulfide bond ^18^. CD5L is a member of the scavenger receptor cysteine-rich (SRCR) family and consists of three SRCR domains in both murine and human CD5L ^20^. It was originally discovered as Spα or apoptosis inhibitor of macrophages (AIM), being expressed predominantly by macrophages ^21^. In circulation, it has been speculated that CD5L binds to IgM to avoid renal excretion ^22^, whereby the protein is released under certain conditions. However, it is unclear when, where and how frequently CD5L incorporation happens. Disconcertingly, reported circulatory CD5L concentrations differ wildly, ranging from 0.1 to 60 mg/L (i.e., ∼500 fold) ^20, 22–24^, possibly reflecting limitations to detect either the free or putatively IgM-bound form. While free CD5L has been reported to induce a plethora of immunomodulatory effects, the exact functions of this protein remain largely unknown ^25–27^. Above all, the incidence and significance of the IgM-bound form of CD5L are currently undetermined. Here, we characterize human circulatory IgM for its structural composition and association with CD5L. Combining molecular biology and mass spectrometry (MS)-based techniques, we redefine the circulatory IgM complex and explore the role of CD5L in complement activation and Fc receptor binding.

## Results

### CD5L occurs synchronously with IgM in circulation

We started our characterization of IgM and CD5L by tracking the abundances and associations of these proteins in serum from healthy donors (n=42). To investigate the abundance of the individual protein chains (Igµ, J, CD5L) in a way that is unbiased to structure and complex formation, we used bottom-up proteomics with label-free quantitation **(Figure 1A, S1)**. Using this approach, we detected an average total CD5L serum concentration of 1.7 µM or 60 mg/L, consistent with the higher end of literature values ^20^. Remarkably, we found that CD5L levels correlated very tightly with those of the IgM heavy chain constant region (IgµC, R = 0.98), hinting at high levels of circulating IgM-CD5L complexes. Furthermore, the observed CD5L/IgµC molecular ratio was consistently ca. 0.15, translating to close to one CD5L molecule per IgM pentamer.

**Figure 1.**
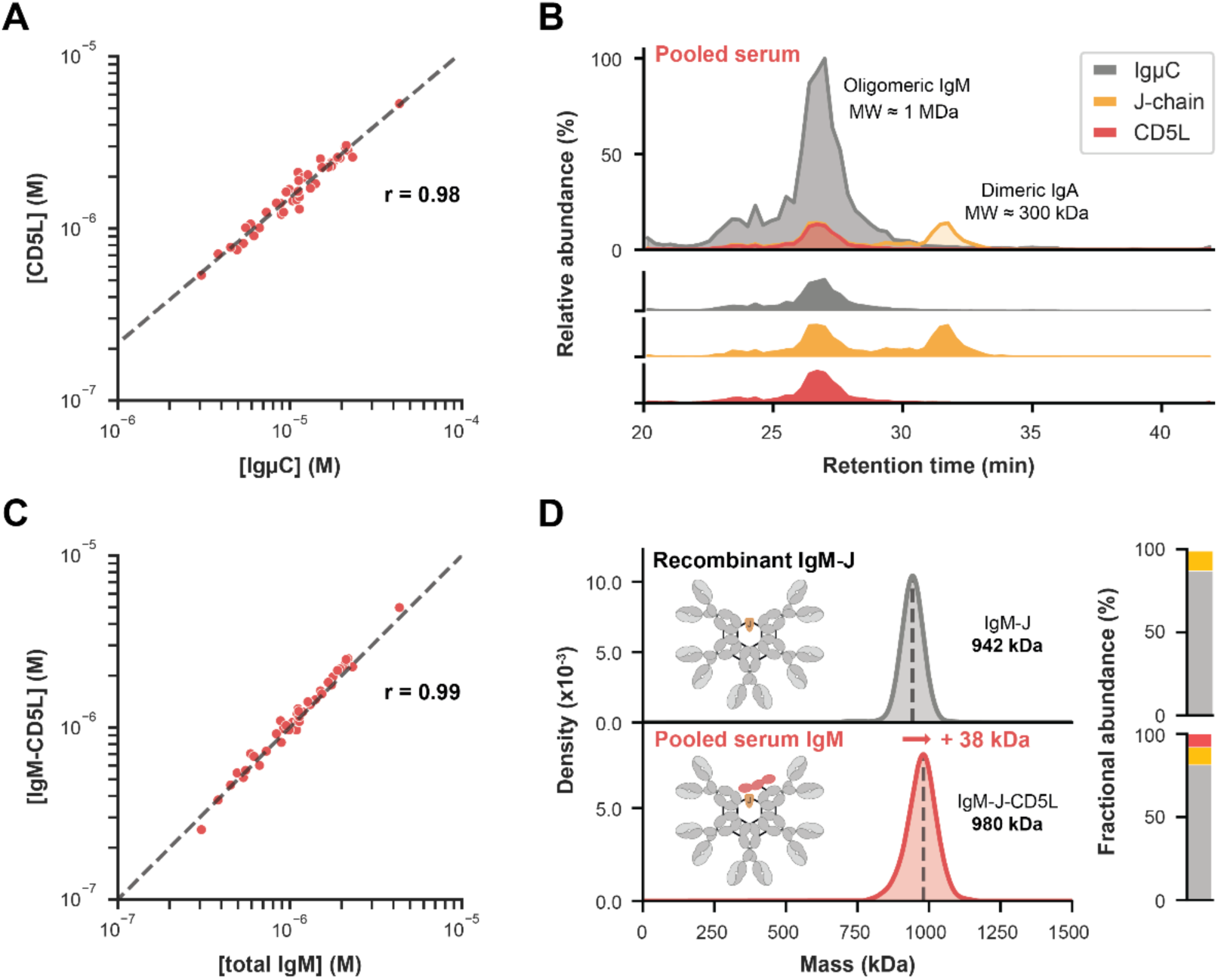
All circulatory IgM is pentameric with J-chain and CD5L incorporated. **A)** Total levels of CD5L and IgM heavy chain (IgµC) show a strong correlation in serum (r=0.98). Absolute concentrations were assessed by label-free quantitation by proteomics. The molecular ratio between CD5L and IgµC was approximately 1:7, advocating the incorporation of one CD5L molecule per IgM pentamer. The dotted line indicates a linear regression model fitted to logarithmically scaled data. **B)** In complex-centric protein profiling of pooled serum, CD5L elutes exclusively (> 99%) with IgµC and J-chain as a single complex of ca. 1 MDa. Shown are the elution profiles of the individual protein chains obtained from proteomics on individual size-exclusion chromatography (SEC) fractions. The secondary elution peak of the J-chain corresponds to incorporation in dimeric IgA. **C)** Levels of IgM-CD5L complexes and total IgM ELISA similarly show a high correlation in serum (R=0.99). The ELISA makes use of a mAb (5B5) that can recognize CD5L when bound to IgM. Quantification was based on a recombinantly produced IgM-J-CD5L complex standard **(Figure 2, S5)**. The ratio between IgM-CD5L complexes and total IgM was approximately 1:1, implying that all serum IgM incorporates a CD5L molecule. The grey line indicates this 1:1 molecular ratio. **D)** The principal configuration of circulatory IgM is a pentamer with J-chain and CD5L. Combining mass measurement by CD-MS (left) with proteomics (right) confirmed that recombinant IgM-J (targeting wall teichoic acid of *S. aureus*) is a pentamer with J-chain ((IgM)5:(J)1) (top). IgM purified from pooled serum was similarly homogeneous (bottom), though the average mass was shifted by +38 kDa and CD5L was detected by proteomics. This matches the mass increase expected for the incorporation of one CD5L molecule to form ((IgM)5:(J)1:(CD5L)1).

To further investigate a putative structural relationship between IgM and CD5L, we next subjected pooled sera to size-exclusion chromatography (SEC) MS. In this approach, bottom-up proteomics was applied to SEC fractions to generate chromatograms of individual proteins, revealing complexation based on synchronous elution. Strikingly, the chromatogram of CD5L revealed essentially complete (> 99%) synchronous elution with IgM as well as with the J-chain in the higher MW fractions around 1 MDa **(Figure 1B)**. No secondary CD5L elution peak was observed in lower MW fractions, indicating that the bulk of CD5L is in association with IgM and the J-chain. Also in SEC MS, we observed an intensity ratio of about 1:1:10 between CD5L, J-chain and IgµC, again hinting towards a 1:1:1 molecular ratio of pentameric IgM, J-chain and CD5L in circulation.

### IgM in circulation is principally a J-chain-linked pentamer with CD5L

Having demonstrated that CD5L is tightly connected to IgM in circulation, we next proceeded to characterize the exact molecular composition of circulatory IgM. First, to directly measure and quantify levels of CD5L-IgM complexes, we set up an ELISA that specifically measures these complexes. For this, we generated a dedicated mAb (5B5) that recognizes CD5L when it is bound to IgM **(Figure S2)**. Using this approach, we found that in the serum of 42 healthy donors, the levels of IgM-bound CD5L and total IgM also showed a near-perfect correlation (R = 0.99) and a molecular ratio of 1 (**Figure 1C**), further implying that IgM exists mainly as a CD5L-containing complex in circulation. In contrast, employing a set-up that solely detects unbound CD5L, we found those levels averaged 0.7 mg/L, thus making up only a small fraction (∼1%) of the total circulating CD5L population (**Figure S2D**).

To elucidate the exact molecular composition of circulatory IgM, we next subjected IgM purified from pooled serum and the plasma of individual donors to two single-particle mass measurement techniques, native charge detection (CD) MS ^28^ and mass photometry ^29^. Combining these precise intact molecular weight measurements with bottom-up proteomics to identify constituent proteins, allowed us to establish the exact stoichiometries and possible co-occurrences of different oligomeric states (**Figure 1D** and **S3**). For comparison we also analyzed a recombinant IgM-J and observed it to be exclusively a J-chain-linked pentamer with a measured mass of 942 kDa ((IgM)_5_:(J)_1_). For each serum or plasma IgM sample, we consistently observed a mass of about 40 kDa higher, a shift that closely matches the incorporation of one 36 kDa CD5L molecule. Simultaneously, proteomics analyses on the same circulatory IgM samples consistently revealed the presence of CD5L at a molecular ratio of about 1:1:10 (CD5L:J-chain:IgµC). Notably, circulatory IgM was also highly homogeneous in mass and therefore oligomeric state, without evidence for the existence of IgM hexamers in any of the samples analyzed (abundance < 1%) **(Figure 1)**. We thus conclude that the canonical form of human circulatory IgM is a J-chain-linked pentamer with one CD5L molecule ((IgM)_5_:(J)_1_:(CD5L)_1_).

#### CD5L is not expressed by B cells and is covalently integrated into IgM via CD5L-Cys191

The near-complete integration of CD5L into IgM as observed here would be best explained by secretion from B cells as a fully formed complex. However, the expression of CD5L is reported to be mostly restricted to macrophages ^21, 30^. To investigate the possibility of co-expression of CD5L and IgM in B cells, we cultured naïve and memory B cells, both in a T cell-dependent and T cell-independent manner and analyzed IgM in the supernatants **(Figure 2A)**. The secreted IgM was completely devoid of CD5L, demonstrating that B cells do not produce CD5L. Recombinant co-transfection of IgM-J and CD5L within the same cell proved to be challenging and did not lead to efficient complex formation. In contrast, co-culture of IgM-J-expressing and CD5L-expressing cells reliably produced homogenous IgM-J-CD5L (**Figure S4**). This implies that complex formation of B cell-derived IgM with CD5L occurs in the extracellular space after both IgM-J and CD5L have been expressed independently.

**Figure 2.**
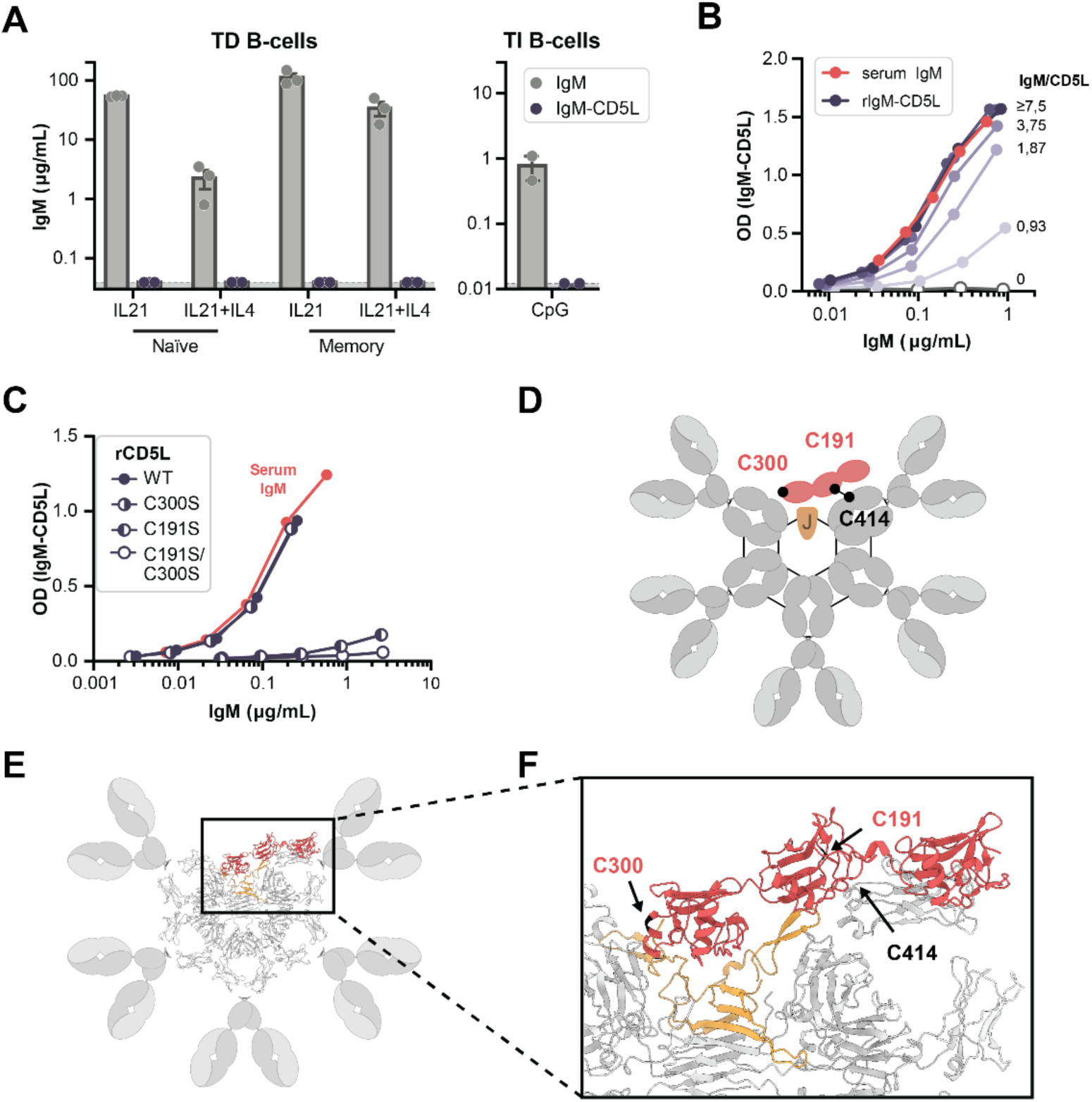
Formation of IgM-CD5L complexes. **A)** Isolated human peripheral blood B cells were cultured *in vitro* with both T-dependent (TD) and independent (TI) stimuli, after which supernatants were analyzed for secreted IgM and IgM-CD5L complexes by ELISA. Only IgM devoid of CD5L was detected, irrespective of culture conditions. Data shown are from 2 and 3 different donors. **B)** *In vitro* generation of recombinant IgM-CD5L complexes. Different molar ratios of CD5L to IgM (clone 2D5 ^14^) were incubated in the presence of 0.1 mM GSH/GSSH. Complex formation was assessed by ELISA. With a molar excess of CD5L, saturation was observed. Representative plots of n=2 experiments. **C)** Human CD5L contains two predicted unpaired cysteines, C191 and C300, that could potentially interact with IgM. Recombinant CD5L was produced with C191S or C300S mutations or both and was tested for their ability to form complexes with IgM as determined by ELISA. C191S disrupts complex formation, whereas C300S does not. Representative plots of n=2 experiments. **D)** Schematic representation of IgM-CD5L complex with highlighted C191 and C300 of CD5L and C414 of IgM-Fc. **E)** Proposed structural model of IgM-CD5L complex. AlphaFold2 model of CD5L and J-chain was fitted into the structure of IgM core/Fc region with J-chain (PDB: 8ADY). **F)** Detailed view on CD5L and J-chain within IgM.

To further clarify the binding mechanism of CD5L to IgM, we recombinantly produced CD5L (rCD5L) and pentameric IgM-J and studied CD5L incorporation *in vitro* **(Figure 2B** and **S5)**. We observed an efficient association of rCD5L with IgM under mild reducing conditions, implicating disulfide bond formation between IgM and CD5L to be an integral part of complex formation.

Saturation of IgM binding was reached upon incubation with an excess of CD5L. These binding experiments were repeated with CD5L purified from serum IgM (sIgM-CD5L) and with recombinant IgM-Fc, producing similar results **(Figure S5)**. Integral complex formation was confirmed by shielding of CD5L epitopes for two distinct anti-CD5L mAbs (10D11, 7E4) that do not bind serum-derived IgM **(Figure S2E)**.

Based on a structure prediction by Alphafold2 ^31^, human CD5L contains two unpaired cysteines, C191 and C300. To assess their role in IgM-J-CD5L complex formation, we produced recombinant hCD5L variants with these cysteines mutated to serines and tested their ability to associate with IgM-J **(Figure 2C)**. Complex formation was severely reduced by a C191S mutation, but essentially unaffected by a C300S mutation. This suggests that only C191 could be disulfide-linked to IgM. This data is in line with the earlier proposed IgM-CD5L linkage via the homologous C194 in experiments wherein murine IgM was reconstituted *in vitro* with murine CD5L ^18^ (**Figure 2D** and ^18^). To further elucidate the interaction, we next subjected CD5L, J-chain and two IgM-Fc regions to structure prediction by Alphafold2 multimer.^31^ The model positioned CD5L in the gap of the IgM-Fc pentamer but somewhat sticking out of the IgM-Fc plane (**Figure 2E, F**). The SRC3 domain is predicted to be positioned closest to the junction formed by the J-chain, in between the N-and C-terminal loops of the J-chain. The SRC2 domain is located above the cleft between the Fc-Cμ4 and Fc-Cμ3 domains and the N-terminal loop of the J-chain, positioning C191 in close proximity to an unpaired C414 of IgM. The SRC1 domain is expected to be flexible. Taken together, CD5L thus binds the gap of IgM-J and involves disulfide bond formation requiring C191.

### IgM-J-CD5L equally activates complement but is less potent in pIgR FcαµR binding

We next assessed the putative functional consequences of CD5L incorporation into IgM. First, we investigated a potential effect in the activation of the classical pathway of the complement system, for which IgM can be a potent activator **(Figure 3A** and **S6)**. We used a recombinant monoclonal IgM antibody specific for biotin with and without CD5L and assessed its ability to induce C3b deposition in ELISA and cell lysis of biotinylated human red blood cells ^14^. Both rIgM-J and rIgM-J-CD5L equally induced C3b deposition and cell lysis. We likewise investigated a potential effect of CD5L on C3b deposition on bacteria, using a monoclonal antibody that binds the wall teichoic acid glycopolymer on *S. aureus* ^32^, but again found no differences in activities **(Figure S6D)**.

**Figure 3.**
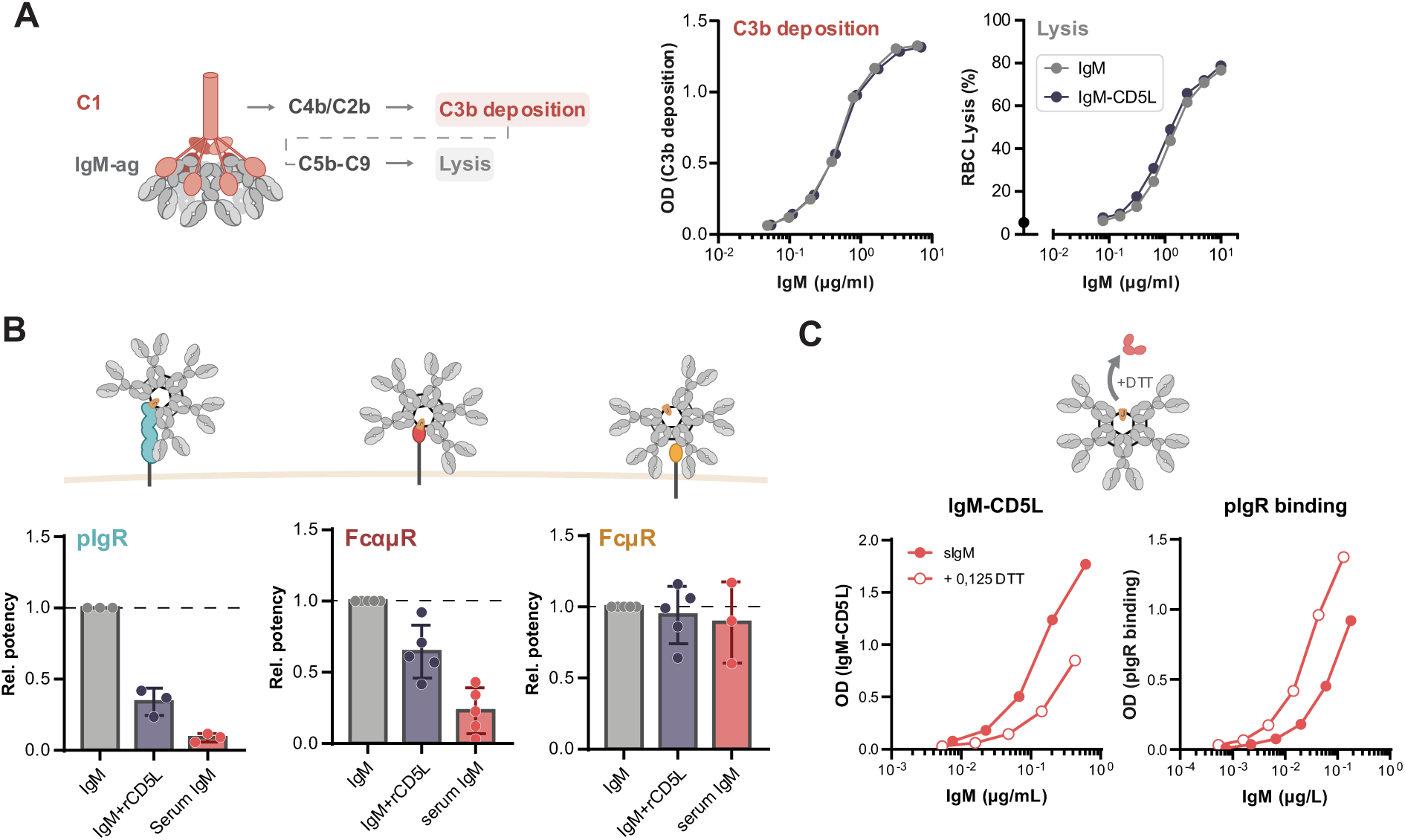
CD5L in IgM decreases pIgR and FcαµR binding; complement activation and FcµR binding unaltered. **A)** Complement activation by IgM was assessed by measuring C3b deposition or red blood cell (RBC) lysis upon binding of recombinant anti-biotin IgM +/-CD5L to plates coated with biotinylated albumin, or biotinylated RBCs, respectively. Representative plots of n=3 experiments. **B)** Binding of recombinant IgM (2D5) +/-CD5L and serum IgM to different IgM-binding receptors. Whereas binding to FcµR is unaffected by CD5L, binding to both pIgR and FcαµR is reduced for CD5L-containing IgM. Data from n=3-5 experiments. **C)** Upon limited reduction of serum IgM, selective release of CD5L proved feasible (left panel) which resulted in increased pIgR binding (right panel). Data representative of n=3 experiments. Bars and error bars represent mean and S.E., respectively.

We next proceeded to characterize the interaction of IgM-J-CD5L with IgM Fc receptor proteins (FcRs) in ELISA. These included pIgR, FcµR and FcαµR, which are involved in mucosal transport ^5^, regulation of lymphocyte responses ^3^, and immune complex uptake by antigen-presenting cells ^4^, respectively. To this end, we measured the binding of recombinant and serum-derived IgM complexes to FcRs **(Figure 3B** and **S7A)**. We found that the binding of rIgM-J-CD5L to both pIgR and FcαµR was reduced compared to rIgM-J, while binding to FcµR was unaffected. Also serum-derived IgM (containing CD5L) displayed reduced binding. Binding to pIgR was also assessed using rIgM-Fc and in a reverse binding experiment, both with results corroborating our initial findings **(Figure S7B)**. To confirm that these effects were caused by CD5L, we also tested whether the release of CD5L from serum IgM would enhance binding to pIgR and FcαµR. Selective release of CD5L from serum IgM upon limited reducing conditions proved feasible **(Figure 3C and S7C)**. This resulted in increased binding to the pIgR and FcαµR, while binding to the FcµR was not affected **(Figure S7D)**. Combined, CD5L incorporation reduces the binding of IgM-J to pIgR and FcαµR, but has no substantial effects on binding to FcµR or complement activation by IgM-J.

### Secretory IgM is principally devoid of CD5L

Secretory IgM is known to be associated with the secretory component (SC), the extracellular portion of the pIgR, which is cleaved from the cells enzymatically after transcytosis to mucosal apical sides. Based on the reduced reactivity of IgM-J-CD5L with pIgR, we hypothesized that, unlike circulatory IgM, secretory IgM may be devoid of CD5L. In stark contrast to serum, we found that CD5L levels in saliva and milk determined by proteomics were much lower and did not correlate with IgµC (**Figure S8**). In agreement, ELISA detection of IgM-bound CD5L in secretory fluids was also much lower than in serum and did not match that of total IgM. Instead, we found a high abundance of SC and a 1:1 molecular ratio between IgM-bound SC and total IgM. In contrast to circulatory IgM, secretory IgM is mostly devoid of CD5L and contains the SC instead.

## Discussion

In this study, we redefined the molecular composition of IgM and revealed its intimate link with the scavenger protein CD5L in circulation. Strikingly, we have now shown that the principal form of circulatory IgM is a J-chain-linked pentamer in complex with CD5L. Conversely, while secretory IgM also consists of J-chain linked pentamer, it is mostly devoid of CD5L but associated with the SC. We speculate that this distinction has eluded discovery due to the relatively small size of CD5L, its assembly after secretion of IgM-J, and the lack of dedicated assays.

While CD5L has been known to co-purify with IgM, its relatively small size (about 4% of the IgM mass) makes it easy to overlook its presence and quantify its abundance. Equally, immunoassays are likely to have underestimated CD5L levels since they were not optimized to include the detection of the IgM-bound form. Nonetheless, several recent plasma proteomics studies revealed high concentrations of total CD5L and a tight correlation with IgM abundance ^33–35^, in line with our data and an earlier study by Arai et al. ^22^. In contrast to the J-chain, CD5L is not B-cell derived and is thus integrated into IgM after its expression by a different cell type. Many studies on antibodies have traditionally been performed using immortalized hybridoma B cells and may have wrongly inferred that the resulting CD5L-devoid IgM molecules are representative of circulatory IgM.

While several impressive high-resolution cryo-EM structures of IgM have been published in recent years ^15–17, 19^, the authentic picture of circulatory IgM is thus still incomplete. Most of these studies used *in vitro*-produced IgM without CD5L, with only one negative-stain EM study describing the putative binding of murine CD5L to murine IgM, when reconstituted *in vitro* ^18^. A very recent cryo-EM study resolved a structure of serum-derived IgM ^17^, but surprisingly, also here, CD5L is lacking. We speculate that this may be due to the use of a myeloma-derived sample in this study, wherein IgM may be overproduced to a level where CD5L incorporation is much lower. Further structural studies on healthy human IgM that does include CD5L are thus highly encouraged.

Also, the role of (free) CD5L may need some rethinking. Both anti-and proinflammatory effects have been attributed to CD5L^25, 26^, but often such experiments were done in the absence of IgM. High concentrations of CD5L may be produced locally within tissues, but once bound to IgM, CD5L likely needs to dissociate in order to exert its functions ^36^. Of note, mice lacking IgM or J-chain show markedly reduced levels of CD5L, emphasizing the tight link between the two ^22^. We hypothesize that dissociation of CD5L from IgM may occur for instance at inflamed sites, possibly through released glutathione or thioredoxin. Indeed, CD5L levels have been reported to be increased in a number of inflammatory disorders ^23, 24, 37–42^. These dynamics may resemble redox-dependent association/dissociation seen for IgG4 Fab arm exchange ^43^. Alternatively, inflammation may induce increased expression of CD5L by macrophages. Still, the findings of these studies need to be rethought based on our finding that CD5L preliminary associates with IgM in circulation.

Several other exciting questions remain to be addressed. In particular, how and where in the body IgM unites with CD5L during homeostasis as well as during an active (humoral) immune response, and how the complex finds its way into circulation or mucosa? Furthermore, it is currently unclear if the clonal repertoire of IgM is skewed with regard to circulatory or secretory IgM, and how that relates to CD5L incorporation. In theory, this might be akin to the distinction seen in circulatory monomeric IgA and secretory dimeric/polymeric IgA which contains J-chain ^44^.

Regardless, functional separation of IgM with or without CD5L in the circulation or mucosa, respectively, might establish as either a function of the observational decreased affinity to pIgR, and/or less time for IgM produced in local mucosal tissues to acquire CD5L. In addition, these molecules are likely to be functionally distinct due to altered FcαµR binding, which is thought to play a role in immune complex uptake and antigen presentation thereby shaping the adaptive immune response. Therefore, it will be highly important that studies towards the role of IgM in modulating immune responses will explicitly take CD5L into consideration. Also, IgM is more and more considered as an alternative format for therapeutic antibodies, for which incorporation of CD5L might be a relevant consideration. We conclude that these results show that the molecular structure of IgM needs to be redrafted (**Figure 4**), either containing CD5L (in circulation) or SC (in mucosa).

**Figure 4.**
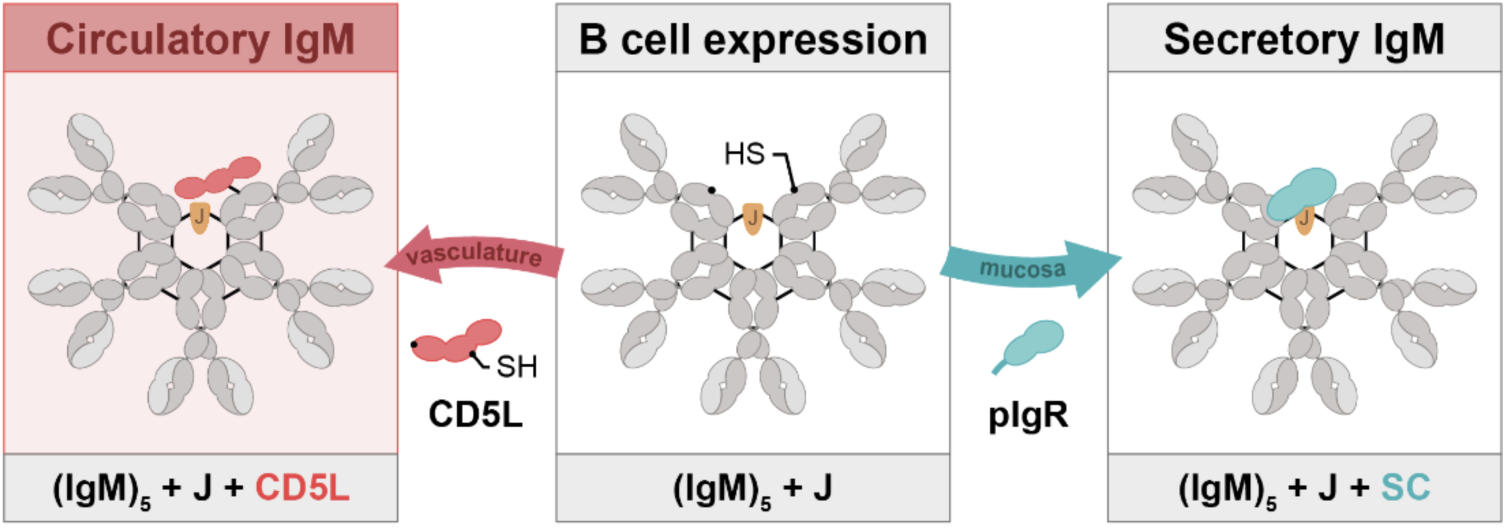
Graphical summary. J-chain coupled IgM pentamers produced in B cells exclusively engages with CD5L when destined to be secreted into the bloodstream, whereas they attach to the secretory component of pIgR *en route* into secretory biofluids such as milk and saliva.

## Acknowledgements

We thank Mads Larsen and Jana Koers for valuable help with the B cell cultures, and René Toes for critical review on the manuscript.

## 1. Supplementary methods

### Serum, saliva, and breastmilk samples

Serum samples were acquired from in-house healthy volunteers who signed written consent or were leftovers from routine diagnostic screening of Tetanus toxoid-boosted donors. Therefore, no informed consent was obtained for the latter samples. Ethical approval was obtained from the Sanquin Ethical Advisory Board. Saliva samples were collected anonymously from healthy volunteers. Human milk was collected by lactating women^8^. Ethical approval was obtained from the Medical Ethics Committee of the Amsterdam UMC, location VUmc and written informed consent was obtained from all participants. All materials were used anonymously without any connection to clinical or person-specific data.

### Bottom-up proteomics sample preparation

All samples, including purified human IgM ±CD5L, as well as whole serum (1 µL), saliva (10 µL), or milk (6 µL) were diluted in a buffer 1% (w/v) sodium deoxycholate (SDC), 10 mM tris(2-carboxyethyl)phosphine hydrochloride (TCEP), 40 mM chloroacetamide (CAA), and 100 mM Tris-HCl at pH 8.0 supplemented with protease inhibitor (cOmplete mini EDTA-free, Roche). Samples were subjected to denaturation and alkylation at 95 °C for 10 minutes, followed by 4-fold dilution with 50 mM Tris-HCl pH 8.5. Proteases were added at an enzyme-to-protein ratio (w/w) of 1:50 for trypsin (Sigma-Aldrich) and 1:75 for Lys-C (Wako), followed by overnight digestion at 37°C. Samples were acidified to roughly 1% formic acid (v/v) to precipitate the SDC, followed by centrifugation cycles for 10 minutes at 20,000 g whereby the supernatant was collected. Samples were stored at -20 °C prior to LC-MS analysis.

### Proteome profiling of human biofluids

Samples were loaded onto Evosep® Pure tips, followed by the regular Evosep protocol except for the washing step, which was substituted by two rounds of 50 µL 0.1% formic acid. Chromatographic separation was performed using the 30 SPD method on an Evosep (Evosep) fitted with an EV-1109 analytical column (Dr Maisch C18 AQ, 1.5µm beads, 150 µm ID, 8 cm length, Evosep), using MilliQ water with 0.1% formic acid (v/v) as solvent A and acetonitrile with 0.1% formic acid (v/v) as solvent B. Mass spectrometric analysis was performed on an Orbitrap Exploris 480 mass spectrometer (Thermo Scientific) using a data-independent (DIA) acquisition method. In each cycle, first, an MS1 scan was acquired at a set resolution of 60,000, followed by 40 sequential quadrupole isolation windows of 15 m/z covering m/z 400-1,000 for HCD MS2 at a set resolution of 15,000. The automatic gain control (AGC) target was set to 300% for MS1 and 1000% for MS2. The maximum injection time was set to 120 ms for MS1 and to “Auto” for MS2. Raw files were searched using DIA-NN (version 1.8) without spectral libraries and with the “Deep learning” option enabled ^1^. Trypsin was set as the protease with a maximum of 1 missed cleavage enabled. Cysteine carbamidomethylation was set as a fixed modification, and N-terminal methionine excision was enabled. The precursor and protein false discovery rates were set to 1%. Normalization was performed per sample type (Milk, Serum, Saliva) using the MaxLFQ algorithm incorporated into DIA-NN. All other settings were set to their default. No imputation was performed.

### Proteomics analysis of purified human proteins

Peptide samples of about 100 ng were separated and analyzed by an UltiMate 3000 UHPLC system (ThermoFisher Scientific) coupled to an Orbitrap Exploris 480 mass spectrometer (Thermo Fisher Scientific). Peptides were first trapped on a 300 μm x 5 mm trap column packed with C18 PepMap100, 5 μm, 100 Å (Thermo Fisher Scientific, P/N 160454) and then separated on a 75 μm × 500 mm analytical column packed with Poroshell 120 EC-C18, 2.7 μm (ZORBAX Chromatographic Packing, Agilent Technologies). Mobile phase A consisted of 0.1% FA (v/v) in MilliQ water, while mobile phase B consisted of 0.1% FA (v/v) in acetonitrile. Peptides were separated at a 300 nL/min flow rate during a 55 min method as follows: 1 min 9% solvent B; a 1 min ramp from 9% to 13% solvent B; a 35 min separation gradient from 13% to 44% solvent B; a 3 min ramp from 44% to 95% solvent B; 4 min 95% solvent B; a 1 min ramp from 95% to 9% solvent B; 10 min 9% solvent B. Mass spectrometric analysis was performed using a data-dependent acquisition (DDA) method. In each cycle, an MS1 scan was required at a set resolution of 60,000, followed by an HCD MS2 scan at a set resolution of 15,000, both using the “Standard” AGC target and “Auto” maximum injection time settings. Data were searched against the UniProtKB/Swiss-Prot human proteome sequence database with MaxQuant version 1.6.17.0 ^2^ using standard settings.

After filtering for proteins with a Q-value ≤ 0.02 and a Score ≥ 50, label-free quantitation (LFQ) values were used to calculate relative abundances.

### Complex-centric proteome profiling by SEC-MS

Complex-centric proteome profiling was performed through proteomics analyses on plasma or milk fractions separated by size-exclusion chromatography (SEC). Chromatography was performed using an Agilent 1290 Infinity HPLC system (Agilent Technologies) consisting of a vacuum degasser, refrigerated autosampler with a 100 µL injector loop, binary pump, thermostated two-column compartment, refrigerated fraction collection module, and multi-wavelength detector. Samples were separated using a dual-column setup comprised of a Yarra SEC-4000 and SEC-3000 column (300 x 7.8 mm i.d., 3 µm, 500 Å or 290 Å respectively) (Phenomenex). Separation was performed at 17 °C with a flow rate of 0.5 mL/min using 150 mM aqueous ammonium acetate with 50 mM L-arginine at pH 7.5 as the mobile phase. A volume of 20 µL pooled serum (ca. 1.2 mg of protein) was injected, followed by separation in 60 min, whereby proteins eluted in a 20-42 min window and were collected in 74 fractions. These fractions were subjected to in-solution digestion and proteomics analysis as described above.

### Production of recombinant proteins

Recombinant IgM-Fc, IgM anti-biotin,anti-CCP (2D5) and anti-WTA were produced as described previously ^3, 4^. To produce recombinant human CD5L, FcµR, FcαµR and pIgR, DNA3.1+ expression vectors containing the complete protein sequence (CD5L) or sequence for the extracellular domains (receptors) were designed. All vectors were created with a C-terminal BirA tag (sequence: GLNDIFEAQKIEW ^5^) and 10xHis-tag, with the exception of the CD5L vector that was tagged N-terminally instead. The vectors were transfected into HEK 293F cells using PEI-MAX according to the manufacturer’s instructions (Invitrogen). After being cultured for 5 days at 37°C, 8% CO2 and shaking at 125 RPM, the supernatant was harvested and filtered through a filter with a pore size of 0.20 µm (Whatman Puradisc 30; Sigma-Aldrich). The recombinant proteins were purified using a HisTrap™ column (Cytiva). The culture supernatant was first dialyzed against phosphate-buffered saline (PBS) and subsequently against a binding buffer (20 mM trisodium phosphate, 0.5 mM sodium chloride, 30 mM imidazole, pH 7.8). The column was washed with water and then binding buffer, after which the supernatant was loaded onto the column. Unbound protein was washed away with binding buffer. The bound proteins were eluted with a gradient of elution buffer (20 mM trisodium phosphate, 0.5 mM sodium chloride, 500 mM imidazole, pH 7.4) and subsequently rebuffered to PBS using a 10 kDa spin column (Amicon Utra-4 Centrifugal Filter Unit; Merck). The concentrations of the purified proteins were determined by measuring the absorbance at 280 nm (NanoDrop One; Thermo Fischer Scientific) after which the samples were stored at -20°C. The purified recombinant proteins were visualized on SDS page using 4-12% Bis-Tris Protein Gels (NuPAGE™; Invitrogen) according to the manufacturer’s protocol, after which they were stained with coomassie (InstantBlue; Expedeon) for at least 30 min or used for western blot. Proteins were site-specifically biotinylated using the Enzymatic Protein Biotinylation Kit (Sigma Aldrich) according to protocol.

Alternatively, IgM-J-CD5L complexes were produced by co-cultures of HEK 293F cells transiently expressing anti-WTA IgM-J or CD5L. Anti-WTA (4497). IgM-J was transfected as described previously ^4^, whereas separately cells were transfected with pcDNA34 expression vectors encoding CD5L under control of the UPE (Cystatin-S) promoter. For the latter, 0.5 µg total plasmid (50% CD5L plasmid/50% empty vector) was transfected per mL of cell culture. After 5h of incubation, IgM-transfected and CD5L-transfected cells were combined in a 1 to 1 ratio and 1mM valproic acid was added to the culture. The produced IgM was purified as described previously ^4^.

### Affinity purification of serum IgM and CD5L

Serum or plasma was diluted 1:1 with PBS supplemented with 500 mM NaCl (mobile phase A), after which samples were loaded on a 10 mL HiTrap column (Cytiva) filled with POROS CaptureSelect IgM affinity matrix (Thermo Fisher) at a flow rate of 1 mL/min using an ÄKTA pure™ 25 (Cytiva). The column was washed with mobile phase A until UV absorption was stable, followed by elution using 100 mM glycine with 500 mM NaCl at pH 3.0 (mobile phase B). Samples were dialyzed overnight against mobile phase A or 20 mM NaAc, 300 mM NaCl (pH 5.5), after which protein concentrations were determined using a NanoDrop (Thermo Fisher Scientific). Samples were stored at -20 °C or -80 °C before further use. Between purifications, the column was cleaned with both 50 mM citric acid pH 2.0 and 6 M guanidine to prevent carryover. Serum CD5L was obtained by reduction of purified serum IgM, with which it co-purifies. Serum IgM was reduced with 1 mM Dithiothreitol (DTT; Calbiogen) for 2h at 37°C, followed by size-exclusion chromatographic separation using a Superdex 200 column (10/300 GL; Cytiva). The fractions containing free CD5L were pooled and dialyzed against PBS and stored at -20°C.

### Generation of monoclonal antibodies specific for CD5L

Monoclonal antibodies against CD5L were obtained as described previously ^6^. In short, a rabbit was immunized with recombinant CD5L and boosted in 4-week intervals. Whole blood was obtained 9 days after the third booster, from which PBMCs were isolated and serum was tested for the presence of CD5L antibodies. B cells were isolated from PBMC samples by FACS sorting. Cells were stained with FITC-labeled mouse anti-rabbit IgG (2A9, Abcam) and with CD5L-bt followed by streptavidin-APC, after which the double-positive were sorted and seeded in single wells. These cells were cultured for 9 days, after which the supernatants were tested by ELISA for specific antibodies against free and IgM-bound CD5L.

RNA was isolated of several positive clones, after which cDNA was synthesized, and IGHV and IGLV regions were amplified using PCR^6^. PCR products were sequenced by Sanger sequencing. Antibodies were expressed using synthetic DNA vectors (Invitrogen) coding the variable regions of the antibodies in combination with mouse IgG1 kappa constant domains, essentially as described before ^6^. Antibodies were produced by transient transfection using the Freestyle HEK293 system, and antibodies were purified as described previously ^3^ using a HiTrap protG column (GE Healthcare) and stored in 5 mM NaAc (pH 4.5).

### Mass photometry

Mass photometry (MP) experiments were performed on a Refeyn One^MP^ or Samux^MP^ mass photometer (Refeyn). Microscope coverslips (24 mm × 50 mm; Marienfeld) were cleaned by sonication in two sequential iterations between isopropanol and MilliQ water, followed by placement of a CultureWell gasket (Grace Biolabs). Typically, 15 µL of PBS was placed in a well for focusing, after which about 3 µL of diluted sample was introduced and mixed. For IgM, the measurement concentration was typically around 20 nM of monomeric subunits. Measurements were recorded for 120 s using medium field-of-view settings on the One^MP^ or standard settings on the Samux^MP^. Calibration of the One^MP^ was performed using an in-house mix consisting of IgG4Δhinge-L368A, IgG1-Campath, apoferritin, and GroEL (73, 149, 479, and 800 kDa). Calibration of the Samux^MP^ was performed using thyroglobulin oligomers (670, 1340, and 2010 kDa). Data were processed in DiscoverMP (Refeyn), followed by analysis and plotting using an in-house Python library.

### Native Orbitrap-based charge detection mass spectrometry

Orbitrap-based CD-MS was performed on an Orbitrap Q Exactive Plus UHMR instrument (Thermo Fisher Scientific). Firstly, proteins were buffer exchanged to 150 mM aqueous ammonium acetate pH 7.5 through six consecutive dilution and concentration steps at 4°C using Amicon Ultra centrifugal filters with a 10kDa molecular weight cutoff (Merck). For experiments studying binding, complexes were assembled by mixing the subcomponents at the desired molar ratios, followed by incubation at room temperature (RT) for at least 30 min. Samples were further diluted using 150 mM aqueous ammonium acetate pH 7.5 to reach the single particle regime, and immediately loaded into gold-coated borosilicate capillaries (prepared in-house) for direct infusion from a static nano-electrospray ionization source. Measurements were performed at low pressure settings using Xenon as collision gas at a set resolution of 200,000 at 400 *m/z*, corresponding to a transient time of 1024 ms. The number of microscans was set to 1, noise threshold set to 0, and no transient averaging was employed. Ion injection times and transmission parameters were manually optimized for each measurement to maintain single ion detection. After acquisition, .raw files were processed and filtered as described by Wörner *et al* ^7^. From the resulting 2-dimensional data, ion events corresponding to species of interest were extracted through density-based spatial clustering of applications with noise (DBSCAN), allowing them to be indicated in a separate color for presentation in mass density figures.

### B cell cultures

B cell cultures were performed analogously as described in ^8^. Buffy coats were obtained from anonymized adult healthy donors upon written informed consent in accordance to the guidelines established by the Sanquin Medical Ethical Committee and in line with the Declaration of Helsinki. Briefly, peripheral blood mononucleated cells were isolated and CD19^+^ cells were isolated. For T cell dependent (TD) conditions, CD27+IgD-(memory) and CD27-IgD+ (naïve) B cell subsets sorted cells were cultured on a monolayer of irradiated 3T3 fibroblast expressing human CD40L ^9^ and in B cell medium supplemented with recombinant human IL-21 (50 ng/mL, Preprotech) or IL-21 and IL-4 (50 ng/mL and 25 ng/mL, Preprotech). For T cell independent (TI) culture, cells were instead supplemented with 0.1 μM CpG (Invivogen). After ten (TD) and seven (TI) days of culture, supernatants were analyzed by ELISA (see below).

### IgM-CD5L complex ELISA

To measure IgM-CD5L complexes, we set up a sandwich ELISA that uses anti-IgM (MH-15-1; Sanquin) for capture, and anti-CD5L (clone 5B5) for detection. 2 µg/mL a-IgM in PBS was coated on maxisorp plates (Thermo Fisher Scientific) overnight at 4°C. After coating, plates were washed five times with PBS supplemented with 0.02% Tween-20 (PBST). Samples were diluted in high-performance ELISA buffer (HPE; Sanquin) to the desired concentration and 100 µL of sample was transferred to each well, after which plates were incubated at RT for an hour while shaking at 300 rpm. A pooled reference serum with a known concentration of IgM was used as a calibrator. After sample incubation, the plates were washed and 100 µL of a-CD5L 5B5-bt diluted in HPE (0.5 µg/mL) was transferred to each well. Clone 10D11 or 7E4 were used in a similar setup to check for proper integration of CD5L into recombinant IgM, as these bind to an epitope on CD5L that is shielded when bound to IgM. Plates were then incubated for 1 h at RT while shaking and then washed again. Subsequently, bound a-CD5L was detected with streptavidin-HRP in HPE (1:1000). Alternatively, in order to determine the total amount of IgM in the tested samples, the IgM complexes were instead detected with a-IgM-HRP (0.33 µg/mL in HPE). Both conjugates were incubated at RT for 30 min while shaking. The detection was visualized and the absorbance was read at 450 nm and 540 nm for background correction. Where appropriate, recombinant CD5L-IgM complexes were used as a calibrator (see below).

### Free CD5L ELISA

To measure CD5L not in complex with IgM, we set up a sandwich ELISA using anti-CD5L clones 10D11 and 7E4 that both only bind to free CD5L, but recognize different epitopes. 1 µg/mL anti-CD5L 10D11 was coated on maxisorp plates (Thermo Fisher Scientific) overnight at 4°C. Plates were subsequently washed five times with PBST, after which samples (100 µL) were incubated at RT (1 hr, shaking 300 rpm) after being serially diluted HPE. After sample incubation, the plates were washed and 100 µL of a-CD5L 7E4-bt diluted in HPE (0.5 µg/mL) was transferred to each well. Plates were further developed as described above for the complex ELISA. Purified recombinant CD5L was used as a calibrator.

### Formation and characterization of CD5L-IgM complexes

Recombinant IgM-J or IgM-Fc-J was combined with an excess of recombinant or serum-derived CD5L in a buffer of PBS supplemented with 0.01% Tween-20, 0.1 mM glutathione (GSH; Sigma) and 0.1 mM glutathione disulfide (GSSG; Sigma) and incubated at RT overnight. After incubation, the complexes were further diluted in PBST to the desired concentration. Formed complexes were tested in the IgM-CD5L complex ELISA and a purified serum IgM sample was included as a reference. To further characterize the formed CD5L-IgM complexes, 20 µg of sample in PBS was fractionated using HP-SEC Agilent 1260 Infinity II (Agilent Technologies) with a Superose® 6 Increase 10/300 GL Column (GE Healthcare). Elution peaks were measured at 280 nm absorbance and the size of these peaks was estimated using multi-angle light scattering (MALS) as described previously ^3^.

### Receptor binding to IgM-CD5L complexes

To determine the impact of CD5L integration of binding to IgM receptors, the binding of these complexes was assessed in ELISA. To this end, 100 µL of biotinylated recombinant FcµR or FcαµR at 1 µg/mL in PBST was added to streptavidin-coated plates (Thermo Fischer Scientific) and incubated for 1h at RT. Plates were washed, after which 100 µL diluted IgM-CD5L complexes diluted in PBS supplemented with 0.2% w/v gelatin (Merck) and 0.1% v/v Tween-20 (PTG) were added to the plate. Plates were incubated for an hour at RT while shaking and then washed. Bound IgM was detected with anti-IgM-HRP (0,33 µg/mL in PTG) for 30 minutes at RT while shaking. Alternatively, for the reversed pIgR binding ELISA, a-IgM was coated at 2 µg/mL in PBS on maxisorp plates, overnight at 4°C. Complexes were added in PTG and incubated for an hour while shaking at RT, after which the plates were washed and biotinylated pIgR (1 µg/mL in PTG) was added to the wells. The plates were incubated at RT for an hour while shaking, then washed and the bound pIgR was detected with HRP-labelled streptavidin (1:1000 in PTG) for 20 min at RT while shaking. Detection was visualized and subsequent measurement of absorbance was performed as described above.

### Complement assays

Complement activation was assessed in a C3b deposition ELISA and complement-dependent cytotoxicity (CDC) assay as described previously ^3^. The C3b deposition ELISA was performed using human serum albumin (HSA; albuman; Sanquin) that was reacted with 30-120 µM biotinylation reagent (EZ-Link™ Sulfo-NHS-LC-Biotin; Thermo Fischer Scientific), whereas for the CDC assay red blood cells were biotinylated at a concentration of 50-500 µM.

For the bacterial killing experiments, *Staphylococcus aureus* Wood46 was cultured from glycerol stock onto Trypticase Soy Agar II with 5% Sheep Blood plates (254087, BD). A single bacterial colony was picked from a plate and grown overnight in Todd Hewitt broth shaking at 37°C. Bacteria were subcultured into fresh Todd Hewitt broth and grown until the mid-log phase (OD600 0.4 – 0.6). The bacteria were pelleted by centrifugation and resuspended in RPMI 1640 medium (Thermofisher/Gibco) with 0.05% human serum albumin (RPMI-HSA) to OD600 = 1.

To quantify antibody binding, bacteria were diluted to OD600 = 0.01 and incubated with a serial dilution of monoclonal anti-WTA (4497) IgM+J ±CD5L antibodies, for 30 minutes at 4°C under shaking conditions. Subsequently, the bacteria were washed with RPMI-HSA and resuspended in 3 µg/mL Goat-anti-human-kappa-AF488 (2060-30, Southern Biotech) for 30 minutes at 4°C under shaking conditions.

To quantify C3b deposition, bacteria were diluted to OD600 = 0.02 and incubated with a serial dilution of monoclonal anti-WTA (4497) IgM+J ±CD5L antibodies for 15 minutes at 4°C under shaking conditions. The bacteria were further diluted to OD600 = 0.01 and incubated with 1% ΔIgG/IgM-serum as a complement source for 30 minutes at 37°C under shaking conditions. ΔIgG/IgM-serum was prepared by depleting human pooled serum from IgG and IgM by affinity chromatography as previously described^10^. After serum incubation, the bacteria were washed with RPMI-HSA, pelleted by centrifugation, and incubated with 3 µg/mL monoclonal mouse anti-C3b (clone bH6), which was randomly labelled with NHS-Alexa Fluor 488 (AF488, Thermo Fisher Scientific) as described previously^11^.

After incubation with detection antibodies, the bacteria were washed, pelleted, and resuspended in RPMI-HSA containing 1% PFA as a fixating agent. Surface binding of fluorescent detection antibodies was analyzed by flow cytometry on a MACSQuant VYB (Miltenyi Biotec).

### Molecular modeling of IgM-CD5L complex

To model the IgM-CD5L complex, AlphaFold2 multimer was used to predict the structure of CD5L along with the J-chain and two IgM-Fc regions ^12^. The CD5L model was then positioned within the previously solved structure of the IgM-Fc pentamer ^13^ by matching the J-chains from both structures, using the Match Maker tool in UCSF Chimera ^14^.

## 2. Supplementary figures

**Figure S1.**
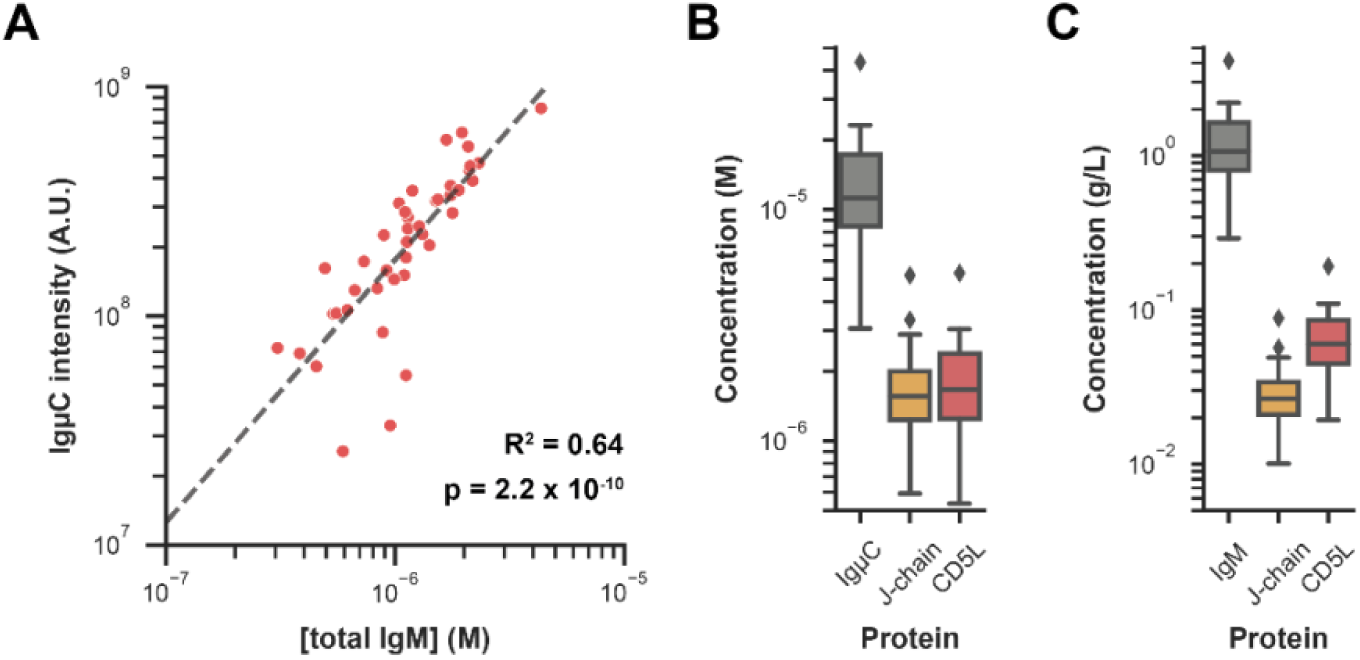
Absolute quantification of IgµC, J-chain and CD5L in serum. **A)** Label-free quantitation intensity values of IgµC correlate well with absolute IgM concentrations determined by ELISA (n=42). The black line indicates a linear regression model fitted to log-transformed data with R^2^ = 0.64 and p = 2.2 x 10^-10^. To estimate absolute concentrations for all detected proteins by proteomics, for each sample, we first calculated an intensity-to-concentration conversion factor using the IgµC intensity and total IgM concentration. This factor assumed that each IgM molecule holds 10 IgµC chains. Then, the intensity values of all other proteins were converted using this conversion factor. **B)** Boxplot showing the resulting molar concentrations of IgµC, J-chain, and CD5L in serum, revealing a close to 10:1:1 molecular ratio. **C)** Boxplot showing the serum concentrations of IgM, J-chain and CD5L in g/L (assuming molecular weights of 950 kDa, 17 kDa and 36 kDa, respectively).

**Figure S2.**
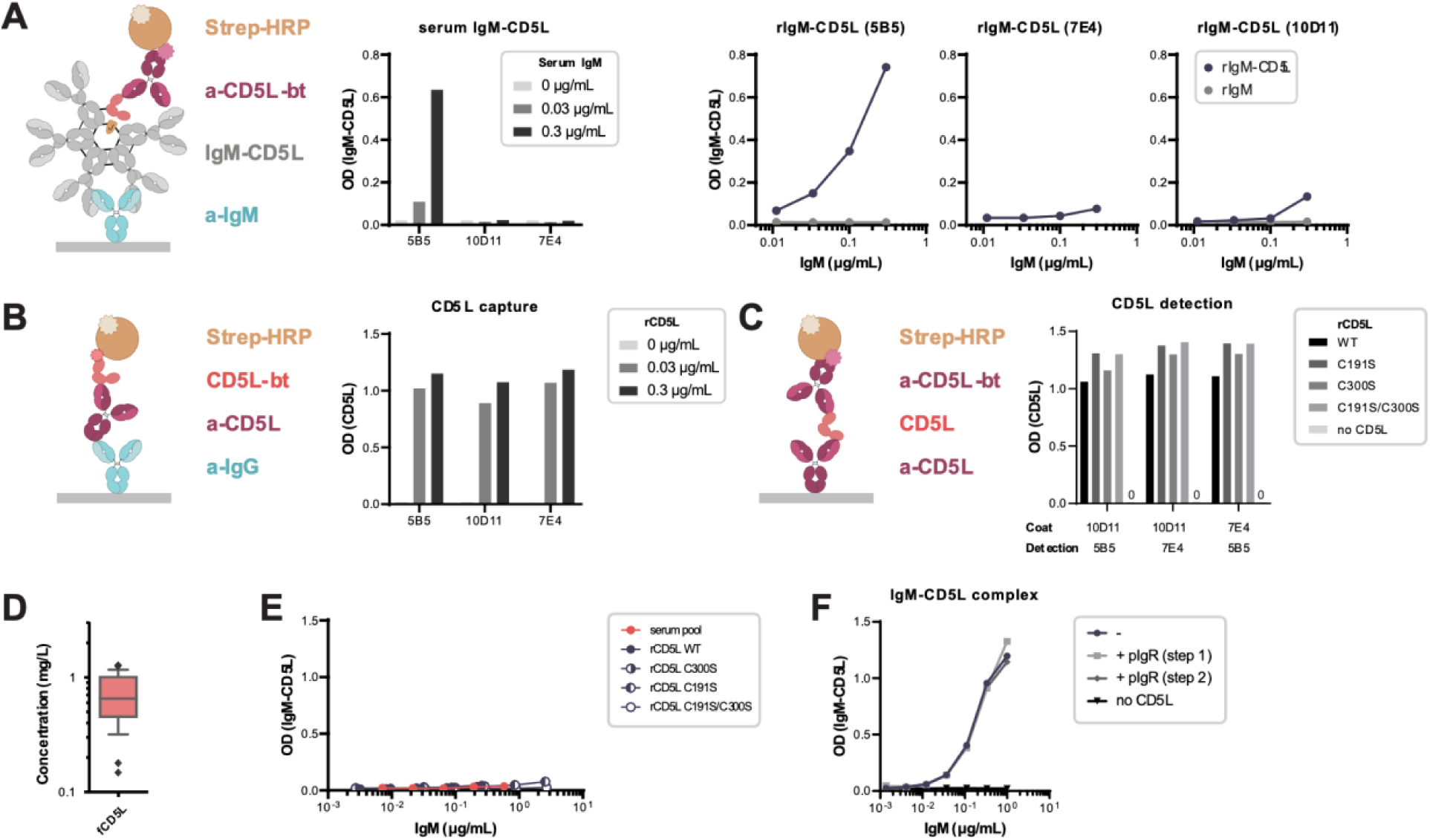
Characterization of monoclonal anti-CD5L antibodies raised to detect IgM-bound or unbound CD5L, respectively. **A)** IgM-bound CD5L was measured in ELISA as depicted, by capture of IgM by an anti-IgM antibody and detection with one of the anti-CD5L mAbs. Only mAb 5B5 was able to detect serum IgM-bound CD5L (left panel). Similarly, 5B5 detects recombinant IgM-CD5L complexes, but does not bind to IgM in absence of CD5L (right panel). The inability of 10D11 and 7E4 to bind to complexes produced *in vitro* indicates that their respective epitopes of CD5L are shielded in these recombinant complexes as for serum IgM-CD5L. **B)** CD5L capture by anti-CD5L mAbs was tested in ELISA as depicted. All clones captured CD5L to a similar extent. **C)** The three anti-CD5L mAbs were tested in combination (i.e. as capture or detection antibody) and bind different epitopes as all combinations gave a signal. Additionally, all clones were able to bind CD5L mutants, showing that detection was not impacted by introduced mutations. **D)** Free CD5L (fCD5L) was measured in serum (n=28) using a combination of 7E4 and 10D11 for capture and detection, which both are unable to bind IgM-bound CD5L. Concentrations were quantified with a standard of recombinant CD5L. **E)** IgM-CD5L complexes formed with WT and recombinant CD5L mutants (Fig. 2C) were also assessed in ELISA with mAbs 10D11 and 7E4 as detection. No CD5L was detected, which implies that CD5L C300S is integrated in a manner that shields similar epitopes as for WT CD5L. **F)** Recombinant pIgR was added either during sample incubation (step 1) or detection (step 2) of the IgM-CD5L complex ELISA with clone 5B5 shown in panel A. The presence of pIgR did not affect the binding of 5B5 to the IgM-CD5L complex, which shows that SC does not interfere with IgM-CD5L measurements and interactions with these antibodies.

**Figure S3.**
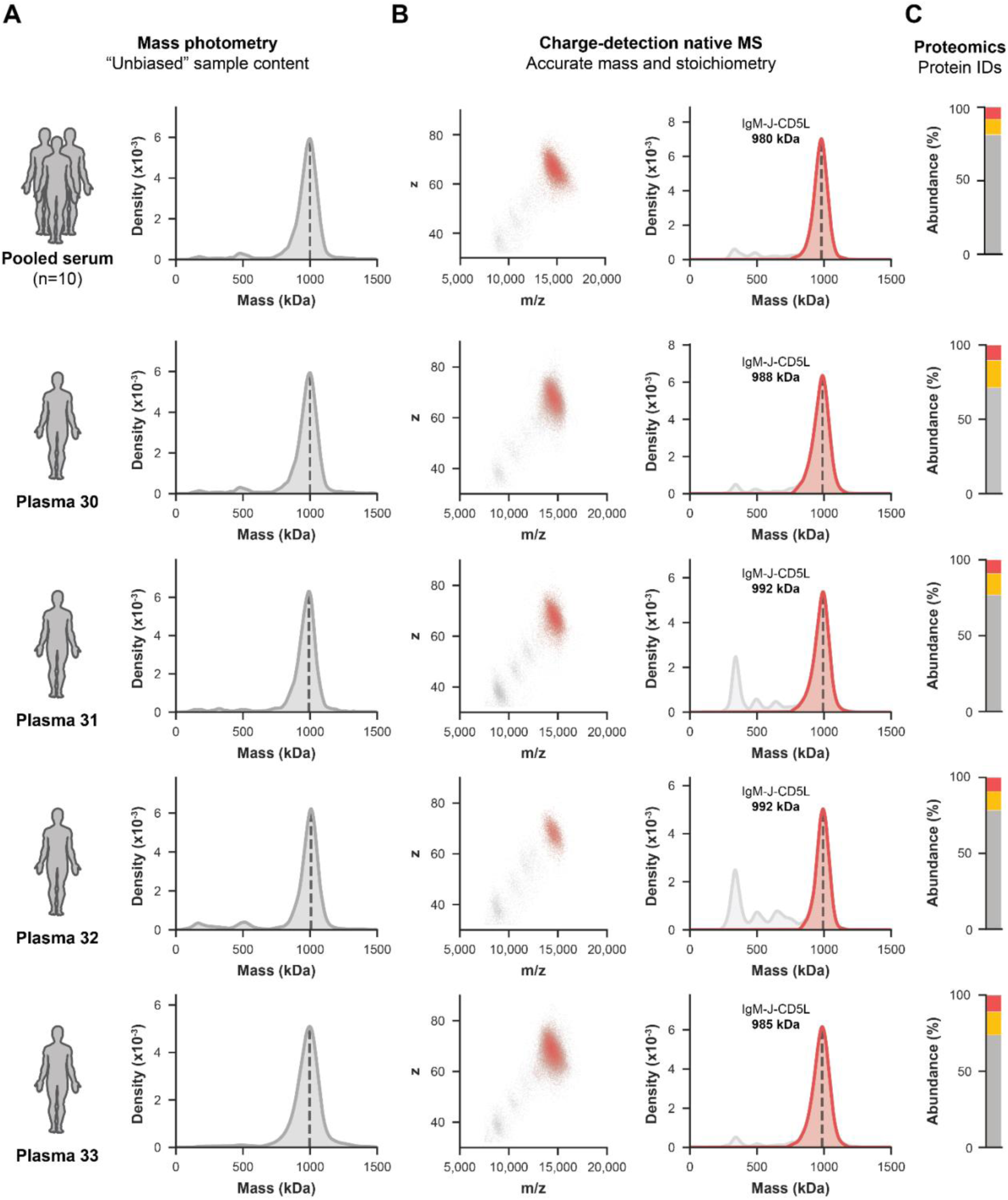
Circulatory IgM from pooled serum and plasma from individual donors is a pentamer with J-chain and CD5L incorporated. **A)** Mass photometry density plots of affinity-purified IgM from pooled serum (n=10, top row) and plasma from four individual donors (bottom 4 rows). All samples were highly homogeneous and showed an average IgM mass of just below 1 MDa. **B)** Analysis of the same samples by CD-MS consistently revealed homogenous mass distributions with average masses between 980-992 kDa. This is a mass shift of +38 kDa compared to a recombinant IgM-J pentamer ((IgM)_5_:(J)_1_, see Figure 1), closely matching the incorporation of one CD5L molecule ((IgM)_5_:(J)_1_:(CD5L)_1_). Shown are the 2D data of *m/z* versus *z* (left), wherein ion events corresponding to IgM were extracted through density-based spatial clustering of applications with noise (DBSCAN), indicated in red. The same measurement is visualized in a 1D mass density plot (right), wherein the extracted IgM ions are overlaid in red against the total measurement in grey. **C)** The presence of CD5L was confirmed by proteomics, with abundances closely matching a molecular ratio of 1:1:10 of CD5L (red), J-chain (yellow) and IgµC (grey) respectively.

**Figure S4.**
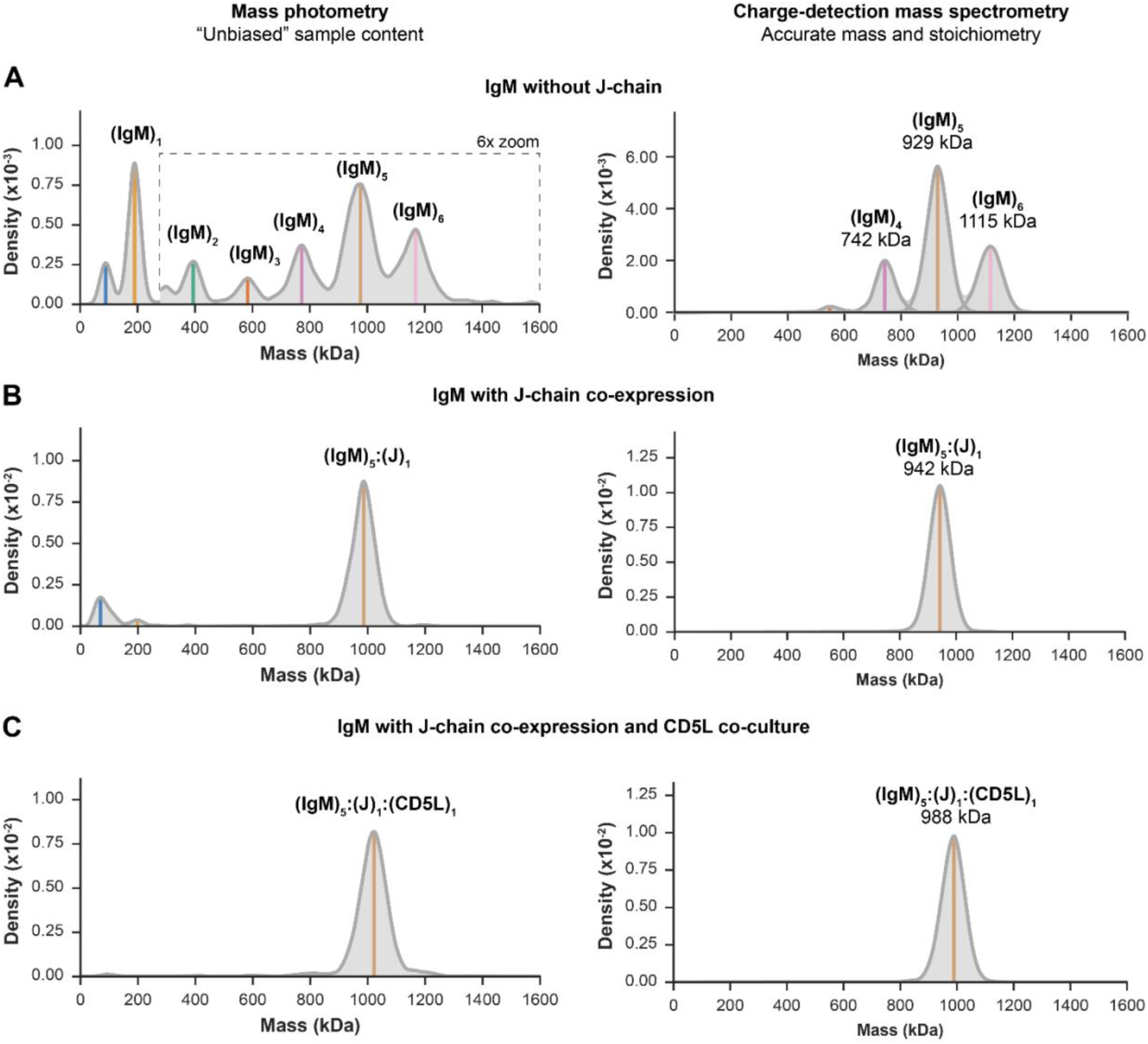
Characterization of recombinant IgM anti-WTA (4497) without J-chain, with J-chain, and with J-chain and CD5L by MP and CD-MS. **A)** IgM without J-chain was produced by co-expression of the HC and LC, resulting in a mixture of oligomers of up to hexamers as measured by MP (left). Accurate mass measurement of larger oligomers by CD-MS confirmed these annotated stoichiometries (right). **B)** Co-expression of IgM with the J-chain resulted in defined oligomerization into exclusively J-chain-linked pentamers. **C)** IgM-J-CD5L was produced by coculture of cells co-expressing IgM and J-chain with cells expressing CD5L, which, after careful balancing, resulted in near-complete CD5L incorporation and stoichiometrically highly homogeneous samples.

**Figure S5.**
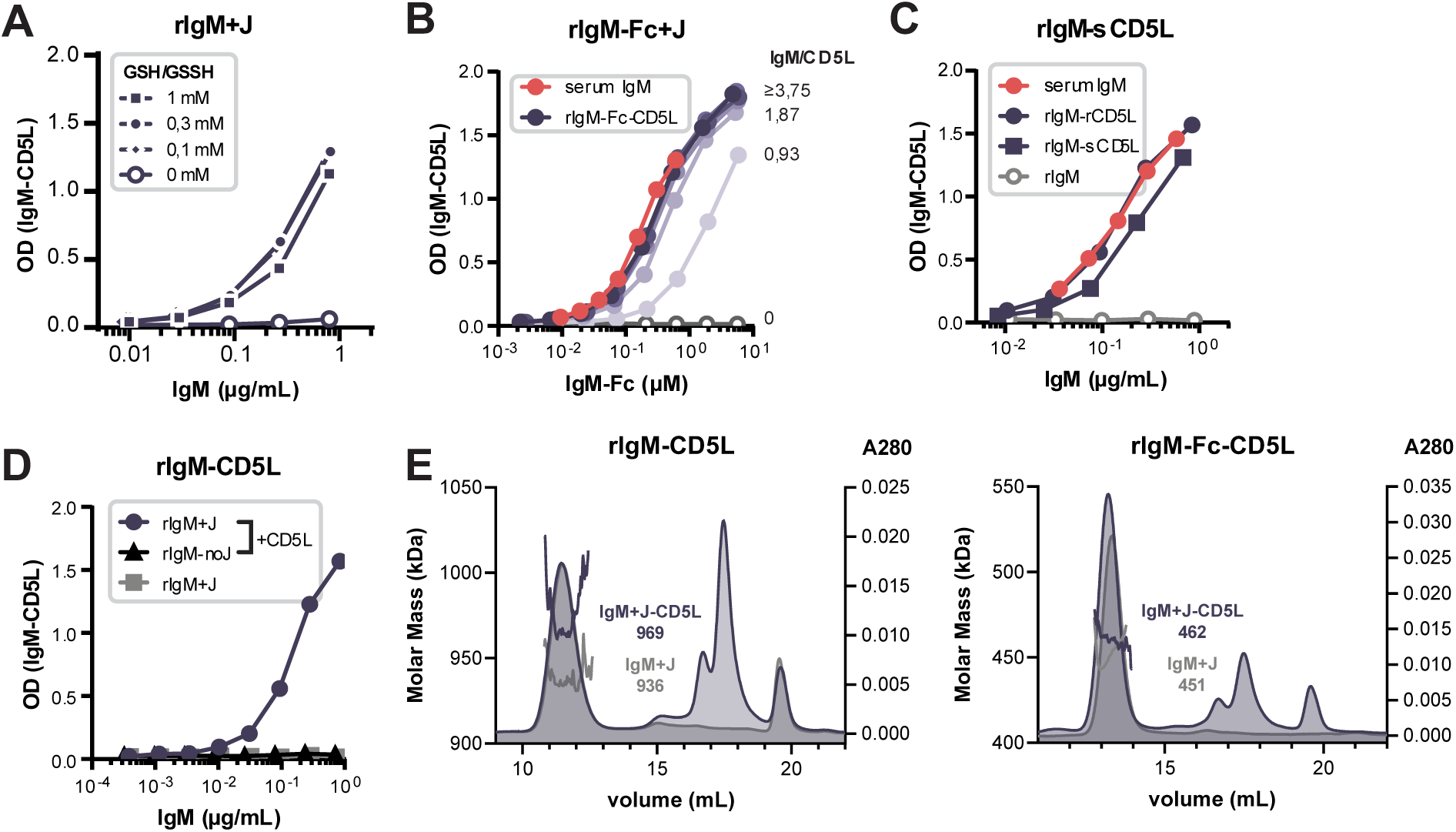
Assessment of IgM-CD5L complex formation *in vitro*. **A)** Recombinant IgM and monomeric free CD5L (molar ratio 1:2) were co-incubated in the presence and absence of a glutathione redox buffer. Only in the latter condition, efficient integration of CD5L into IgM is observed, which implies disulfide bond formation/shuffling is important for complex formation. **B)** Complex formation can also be achieved with recombinant IgM-Fc (CH2-CH4) and CD5L. **C)** Serum CD5L (sCD5L) was purified from serum IgM by reduction with 1 mM DTT for two hours and subsequent HP-SEC, after which the free sCD5L-containing fractions were pooled. sCD5L can, similarly to rCD5L, be (re-)integrated into rIgM. **D)** rCD5L does not bind to recombinant IgM that lacks J-chain and only forms complexes with IgM+J. **E)** Recombinant IgM-CD5L or IgM-Fc-CD5L complexes were assessed by HP-SEC followed by multi-angle light scattering analysis. Elution profiles (right y-axis) and estimated molecular mass (left y-axis) are shown. The addition of CD5L to IgM results in a mass shift corresponding to the integration of one CD5L molecule per IgM.

**Figure S6.**
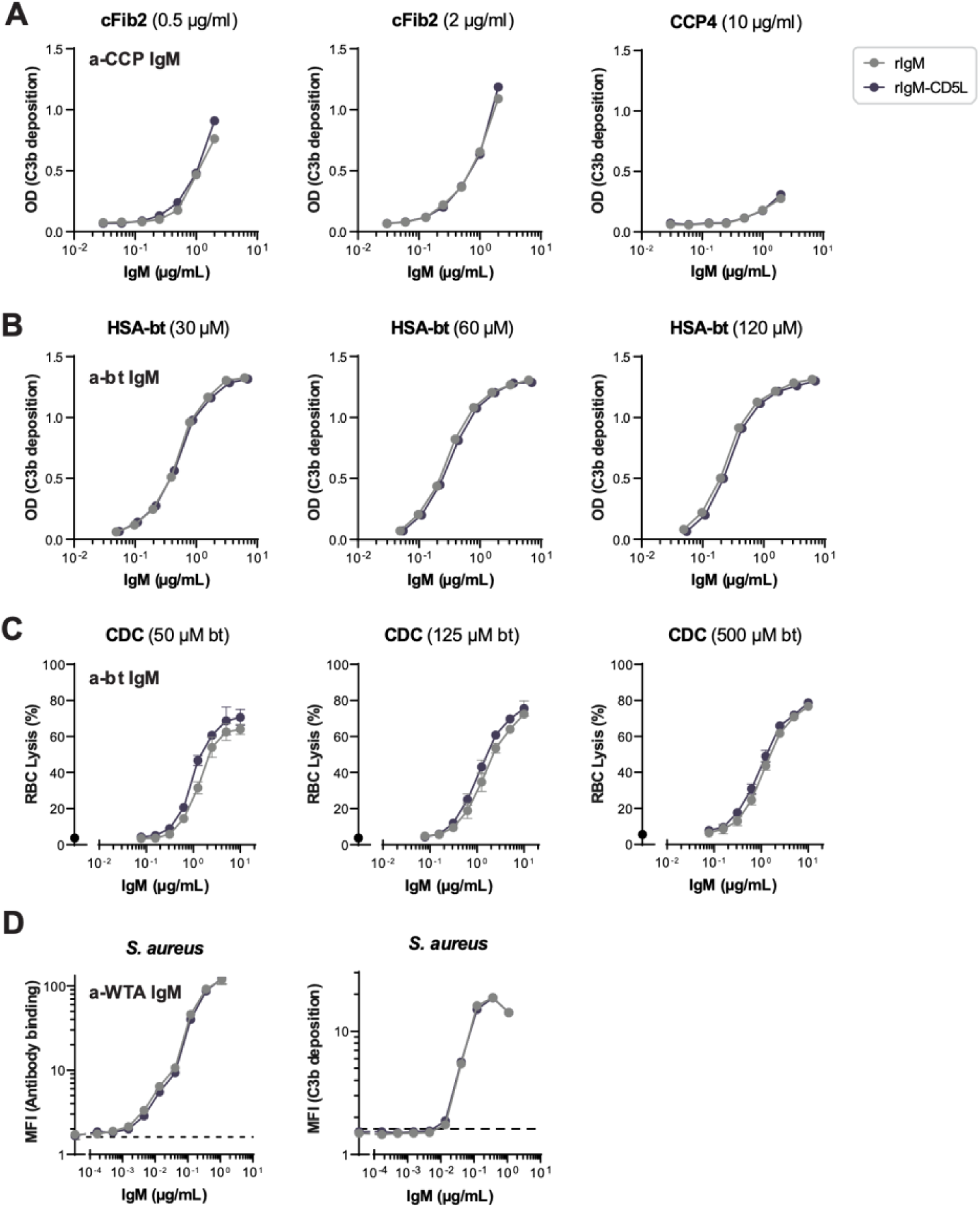
Complement activation by IgM is not affected by CD5L integration. **A)** C3b deposition by an anti-CCP (2D5) or **B)** anti-biotin antibody upon binding to synthetic citrullinated peptides (cFib2 and CCP4) or biotinylated human serum albumin (HSA-bt), respectively, in the presence of 2.5% human serum. Several antigen densities were tested, ranging from relatively low to fully saturated. Representative plots of n=3 experiments **C)** Complement-mediated lysis of biotinylated red blood cells (RBCs) by an anti-biotin antibody in the presence of 10% human serum. RBCs were biotinylated at different concentrations and subsequently incubated with antibody dilutions and serum for 90 min. Released hemoglobin was determined as a measure of cell lysis. Data are plotted as the mean ± SE of n=3. **D)** Antibody binding (left panel) and C3b deposition (right panel) by anti-WTA (4497) antibody on *S. aureus* Wood46. To study C3b deposition, bacteria were opsonized with IgM for 15 min at 4°C and then incubated with 1% human serum depleted of IgG and IgM. Binding of antibodies and C3b deposition were determined by flow cytometry. Dotted line represents the detection of the isotype control. Data are plotted as the mean ± SE of n=3.

**Figure S7.**
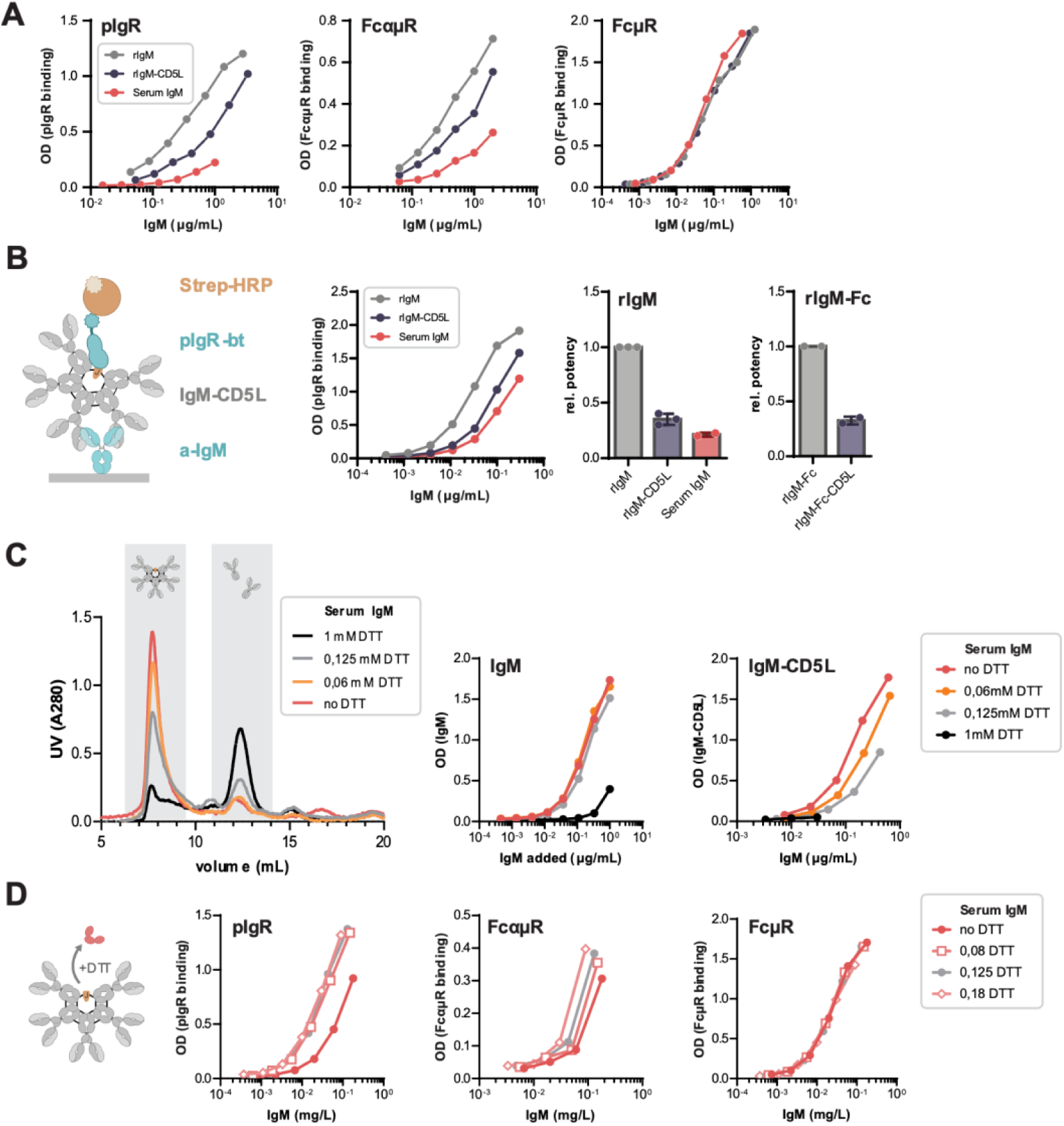
Binding of IgM to pIgR and FcαµR, but not FcµR, is decreased upon CD5L integration. **A)** Representative titration curves for recombinant IgM ± CD5L and serum IgM-CD5L to pIgR, FcαµR and FcµR, of which combined data is shown in Figure 3B. **B)** Binding of IgM and IgM-Fc (CH2-CH4) ± CD5L to pIgR was also assessed in a reversed setup, where IgM was first captured with an anti-IgM antibody and then binding of biotinylated pIgR was detected with streptavidin-HRP. In both approaches, integration of CD5L reduced binding to the receptor. Data from n=2-3 experiments. Bars and error bars represent mean and S.E. **C)** Purified serum IgM was reduced using varying concentrations of DTT ranging from 0.06 mM to 1 mM. Reduced IgM was then assessed using HP-SEC and IgM(-CD5L) ELISA. Whereas at high concentrations of DTT IgM falls apart into fragments the size of IgG-like IgM ‘monomers’, at lower concentrations it remains mostly the size consistent with a pentamer. Nevertheless, at these limiting reducing conditions, CD5L is partially released, as assessed by IgM-CD5L complex ELISA (right panel). **D)** Partial release of CD5L improves binding to pIgR and FcαµR, but not FcµR. Shown is a representative experiment of n=3.

**Figure S8.**
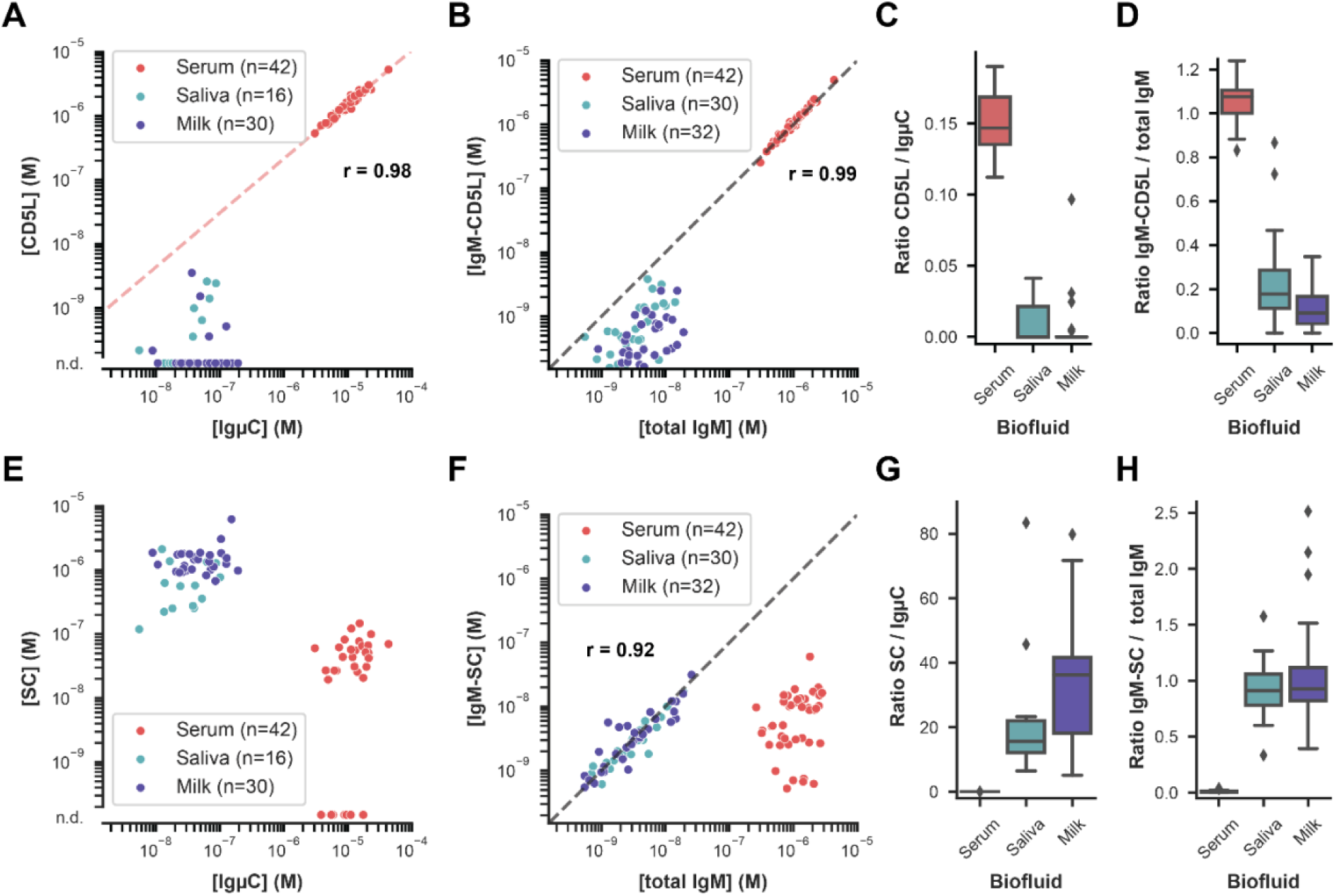
Secretory IgM is largely devoid of CD5L and contains instead the SC of pIgR. **A)** Total levels of CD5L and IgM heavy chain (IgµC) show a remarkably strong correlation in serum (R=0.98) but not in saliva and milk, as determined by label-free proteomics. CD5L levels in saliva and milk were very low, frequently even below the detection limit. These values are indicated at the bottom as not determined (n.d.). The red line indicates a linear regression model fitted to logarithmically scaled serum data. **B)** Levels of IgM-CD5L complexes and total IgM similarly show a high correlation in serum (r=0.99) but not in saliva and milk, as determined by ELISA (see Figure 1). The grey line indicates a 1:1 molecular ratio. **C)** The molecular ratio between CD5L and IgµC was approximately 1:7 in serum, closely matching the incorporation of one CD5L molecule per IgM pentamer. In saliva and milk, this ratio is much lower. **D)** Similarly, while the ratio between IgM-bound CD5L and total IgM was approximately 1:1 in serum, IgM from saliva and milk was nearly devoid of CD5L. **E)** Total levels of SC, the extracellular part of pIgR, were very high in saliva and milk, frequently over an order of magnitude higher than IgµC levels. In serum, SC levels were much lower, sometimes even below the detection limit. These values are indicated as not determined (n.d.). **F)** Consequently, levels of IgM-SC complexes and total IgM ELISA similarly show a good correlation in saliva and milk (R=0.92) but not in serum. The grey line indicates a 1:1 molecular ratio. **G)** The molecular ratio between SC and IgµC was in the order of 10:1 to 40:1 in saliva and milk, highlighting a large excess. In contrast, the ratio in serum was only 0.002:1. **H)** Whereas the ratio between IgM-bound SC and total IgM was approximately 1:1 in saliva in milk, serum IgM was principally devoid of SC.

